# The hidden complexity of Mendelian traits across yeast natural populations

**DOI:** 10.1101/039693

**Authors:** Jing Hou, Anastasie Sigwalt, David Pflieger, Jackson Peter, Jacky de Montigny, Maitreya Dunham, Joseph Schacherer

**Affiliations:** Department of Genetics, Genomics and Microbiology, University of Strasbourg/CNRS UMR 7156, Strasbourg, France; Department of Genome Sciences, University of Washington, Seattle WA, USA

## Abstract

Mendelian traits are considered as the lower end of the complexity spectrum of heritable phenotypes. However, more than a century after the rediscovery of Mendel’s law, the global landscape of monogenic variants as well as their effects and inheritance patterns within natural populations is still not well understood. Using the yeast Saccharomyces cerevisiae, we performed a species-wide survey of Mendelian traits across a large population of isolates. We generated offspring from 41 unique parental pairs, and analyzed 1,105 cross/trait combinations. We found that 8.9% of the cases were Mendelian. Most were caused by common variants showing stable inheritances in a natural population. However, we also found that a rare monogenic variant related to drug resistance displayed a significant and variable expressivity across different genetic backgrounds, leading to modified inheritances ranging from intermediate to high complexities. Our results illustrate for the first time the continuum of the hidden complexity of a monogenic mutation, where genotype is hardly predictive of phenotype.

## Introduction

Elucidating the genetic causes of the astonishing phenotypic diversity observed in natural populations is a major challenge in biology. Within a population, individuals display phenotypic variations in terms of morphology, growth, physiology, behavior, and disease susceptibility. The inheritance patterns of phenotypic traits can be classified as either monogenic or complex. While many traits are complex resulting from variation within multiple genes, their interaction and environmental factors^1^, some traits are primarily monogenic and conform to a simple Mendelian inheritance^2^. Nevertheless, while useful, this overly simplistic dichotomic view could potentially mask the continuous level of the underlying genetic complexity^3-5^. More than a century after the rediscovery of Mendel’s law, we still lack a global overview of the spectrum of genetic complexity of phenotypic variation within any natural population.

Complex traits can be predominantly controlled by variation in a single gene^3^. Similarly, monogenic traits can be influenced by multiple genes in specific genetic backgrounds^5-9^. In fact, it is increasingly evident that monogenic mutations do not always strictly adhere to Mendelian inheritance^6-8^. For example, many human monogenic disorders, including sickle cell anemia and cystic fibrosis, could display significant clinical heterogeneity such as incomplete penetrance and variable levels of severity due to allelic interactions and background specific modifiers^7^. Recent genome-scale surveys of loss-of-function mutations have revealed considerable background effects in various model systems^10-13^ and human cell lines^14-16^, where the mutant phenotypes could be highly variable even between closely related individuals. However, although background effects on “monogenic” loss-of-function mutations are readily seen, such specific mutation type does not reflect the overall genetic diversity and complexity observed in natural populations^17-19^. Specifically, the global landscape of natural genetic variants leading to Mendelian traits has never been thoroughly explored in any species, and their phenotypic effects and inheritance patterns within natural populations are largely unknown.

Here, we carried out a first species-wide identification of causal variants of Mendelian traits in the yeast *S. cerevisiae* to characterize in depth their phenotypic effects and transmission patterns across various genetic backgrounds. We generated a large number of crosses using natural isolates, and analyzed the fitness distribution and segregation patterns in the offspring for more than 1,100 cross/trait combinations. We found that 8.9% of the cases were Mendelian, among which most were caused by common variants and showed stable inheritances across the *S. cerevisiae* species. Interestingly, global phenotypic distribution patterns of multiple Mendelian traits across an extremely large population (∼1,000 isolates) were not necessarily correlated with patterns observed in the offspring from individual crosses. We further characterized a causal variant related to drug resistance and traced its effects across multiple genetic backgrounds. Significant deviations from the Mendelian expectation were observed with variable genetic complexities, illustrating the hidden complexity of a monogenic mutation across a yeast natural population.

## Results

### Global landscape of Mendelian traits in *S. cerevisiae.*

To obtain a comprehensive view of natural genetic variants leading to Mendelian traits in the *S. cerevisiae* species, we selected 41 diverse natural isolates spanning a wide range of ecological (tree exudates, drosophila, fruits, various fermentation and clinical isolates) and geographical sources (Europe, America, Africa and Asia) and performed systematic crosses with one strain Σ1278b (Supplementary Table 1). For each cross, we generated 40 offspring representing 10 individual meiosis (full tetrads), summing up to a panel of 1,640 full meiotic segregants from diverse parental origins (Fig. 1a, panel 1). All segregants as well as the respective parental isolates were tested for 30 stress responsive traits related to various physiological and cellular processes, including different carbon sources, membrane and protein stability, signal transduction, sterol biosynthesis, transcription, translation, as well as osmotic and oxidative stress (Supplementary Table 2). In total, we tested 1,105 cross/trait combinations and analyzed the offspring fitness distribution patterns for each combination (Fig. 1a, panel 2).

**Fig 1.**
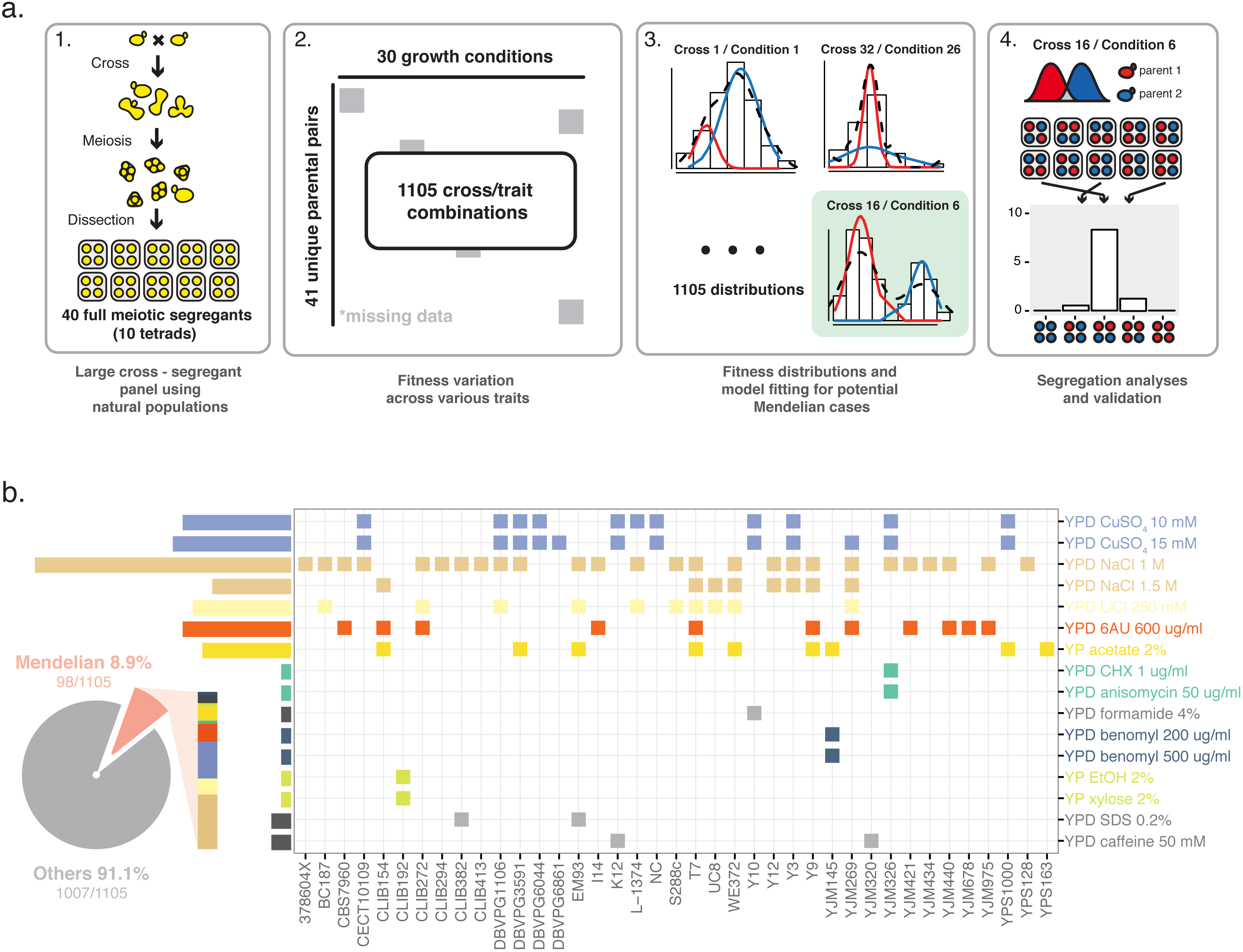
Comprehensive landscape of Mendelian traits in *S. cerevisiae.* (a) Workflow of the detection of Mendelian traits. The workflow was defined as 4 steps, consisting with offspring generation, fitness measurements, model fitting and segregation analysis as indicated. (b) Distribution of all identified Mendelian traits spanning different crosses (x-axis) on conditions tested (y-axis). Each square represents any single Mendelian case and colors indicate different conditions. Pie chart represent the fraction of Mendelian cases relative to the entire dataset.

For a Mendelian trait, contrasting phenotype between the parental isolates was controlled by a single locus, therefore half of the offspring would inherit the causal allele and display a 2:2 segregation in any given tetrads. Consequently, the global offspring fitness distribution would follow a bimodal pattern with equal partitioning of segregants in either parental phenotype cluster. To detect such cases, we first applied a bimodal distribution model with random latent variables for the observed fitness distributions for each cross/trait combination using an Expectation Maximization (EM) algorithm (Fig. 1a, panel 3; Supplementary Fig. 1). A case is considered to fit a bimodal distribution when the observed fitness values could be assigned to two non-overlapping clusters (Fig. 1a, panel 3). For each fitness distribution observed in a given cross/trait combination, the posterior probability that an individual belongs to either fitness cluster was computed (Supplementary Fig. 1), and the general features of the fitted bimodal model such as the means and standard deviations for both clusters as well as their relative ratios were extracted (Supplementary Fig. 2). To determine the cutoff values that allow for high confidence calling of bimodal cases and subsequent cluster assignments, we generated a simulated dataset of 1,000 fitness distributions with the same general features compared to the real data, and reapplied the model fitting procedure (Supplementary Fig. 1 and 2). Using the simulated data as a training set, we determined that a cutoff of posterior probability > 0.8 for cluster assignment while allowing less than 10% of overlapping between the clusters were the best parameters to maintain a high detection performance (area under the ROC = 0.824) while minimizing case loss (Supplementary Fig. 3).

By applying these parameters, 318 cross/trait combinations were detected as bimodal, with the parental isolates belonging to distinct clusters. We then analyzed the phenotypic segregation patterns for all bimodal cases (Fig. 1a, panel 4). In total, 98 cases were identified as Mendelian, displaying the characteristic 2:2 segregation in the tetrads (Fig. 1b). Identified Mendelian cases represented 8.9% (98/1,105) across our sample, and were interspersed among various conditions including large number of instances related to NaCl (28 crosses), CuSO_4_ (13 crosses), 6-azauracil (11 crosses) and acetate (9 crosses) (Fig. 1b). Other low frequency cases were found on conditions related to signal transduction (caffeine), carbon sources (ethanol and xylose) various other conditions (formamide, benomyl and SDS) and the antifungal drugs cycloheximide and anisomycin (Fig. 1b). In addition, we observed co-segregation of unrelated traits (NaCl, acetate and 6-azauracil; Fig. 1b), where the fitness variation patterns in the segregants were highly correlated (Pearson’s correlation ρ > 0.9). We further characterized cases with co-segregations, high frequency cases related to CuSO_4_ and the low frequency case related to resistance to the drugs cycloheximide and anisomycin in detail. For the selected cases, 80 additional full tetrads were tested and the 2:2 phenotypic segregation patterns were confirmed.

### Molecular characterization of identified Mendelian traits

Using bulk segregant analysis followed by whole genome sequencing, we identified one locus for each case as expected. For all crosses displaying co-segregation with NaCl, the same ∼60 kb region (480,000 - 540,000) on chromosome IV was mapped, spanning the *ENA* genes encoding for sodium and/or lithium efflux pumps (Supplementary Fig. 4). While variations of the *ENA* genes were known to lead to osmotic stress tolerance^20^, the phenotypic associations with other co-segregating traits (acetate and 6-azauracil) were previously unknown. Causal genes related to acetate and 6-azauracil were suspected to be in close genetic proximity with the *ENA* locus, however the precise identities of these genes remained unclear. For cases related to CuSO_4_, we mapped a 40 kb region on chromosome VIII (190,000 - 230,000; Supplementary Fig. 4). We identified the *CUP1* gene in this region, which encodes for a copper binding metallothionein (Supplementary Fig. 4). In this case, the common parental strain Σ1278b was resistant to both concentrations of CuSO_4_ tested and the allelic version of *CUP1* in Σ1278b led to stable Mendelian inheritance across multiple genetic backgrounds (Fig. 1b).

Finally, the last characterized case involved two anti-fungal drugs cycloheximide and anisomycin, which was found in the cross between a clinical isolate YJM326 and Σ1278b (Fig. 1b). Pooled segregants belonging to the higher fitness cluster showed allele frequency enrichment for the YJM326 parent across a ∼100 kb region on chromosome VII (420,000 - 520,000; Supplementary Fig. 4). Further analyses yielded *PDR1* as the potential candidate, which encodes for a transcription factor involved in multidrug resistance. Using reciprocal hemizygosity analysis as well as plasmid-based complementation test, we showed that the *PDR1*^*YJM326*^ allele was necessary and sufficient for the observed resistance (Supplementary Fig. 5).

### Fitness distribution of identified Mendelian traits across large natural populations

Although Mendelian traits could exhibit distinctive offspring distribution and segregation patterns in individual crosses, the general phenotypic distribution of such traits within a population was unclear. We measured the fitness distribution of an extremely large collection of ∼1,000 natural isolates of *S. cerevisiae* (The 1002 yeast genomes project - http://1002genomes.u-strasbg.fr/) on selected conditions related to identify Mendelian traits, including resistance to NaCl, LiCl, acetate, 6-azauracil, CuSO_4_ and cycloheximide (Fig. 2). Interestingly, while some traits followed the same bimodal distribution model across the population as was observed in offspring from single crosses (Fig. 2a), other traits with clear Mendelian inheritance pattern in crosses appeared to vary continuously at the population level (Fig. 2b). This observation suggested that the phenotypic distribution within the population might not necessarily reflect the underlying genetic complexity of traits. Instead, the inheritance pattern for any given trait might largely be determined by specific combinations of parental genetic backgrounds.

**Fig 2.**
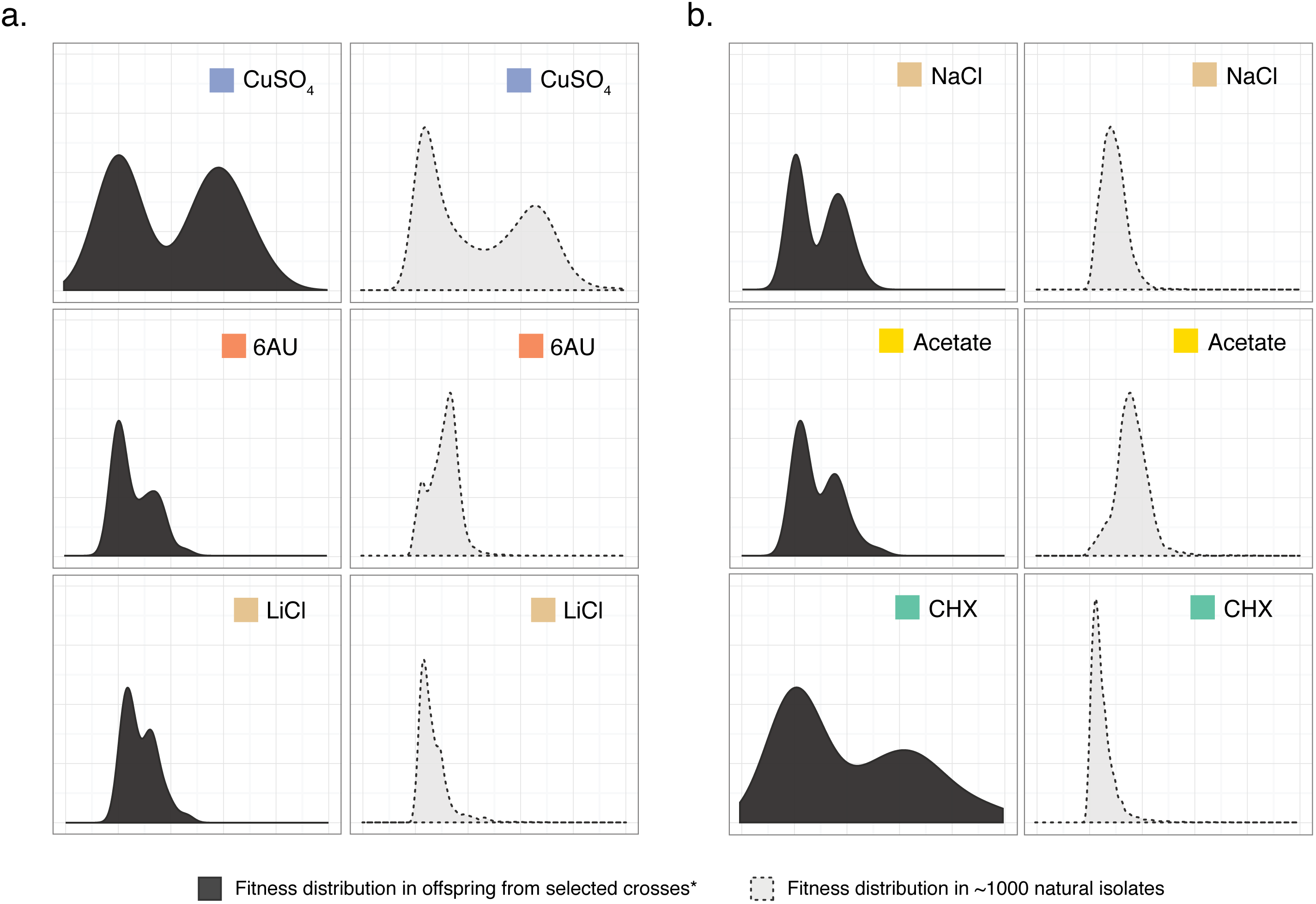
Fitness distribution patterns of identified Mendelian traits within large natural population. Comparisons of the fitness distribution on 6 selected conditions in individual crosses (left panel, N=40) and across ∼1000 natural isolates of *S. cerevisiae* (right panel, N=960) are shown. Conditions tested are color-coded. (a) Bimodal distribution patterns observed both in crosses and at the population level. (b) Bimodal distributions observed only in crosses but not within a population.

### Hidden complexity of a rare Mendelian variant across different genetic backgrounds

While focusing on highly frequent cases such as CuSO_4_ and NaCl provided indications about the transmission stability of common Mendelian variants and revealed previously unknown co-segregations, we were particularly interested in rare cases where the phenotypic effects and the general inheritance patterns across different genetic backgrounds were unknown. The identified Mendelian case related to the anti-fungal drugs cycloheximide and anisomycin could be considered as such. Across our panel, the parent YJM326 was the only highly fit isolate, and few isolates showed similar resistance level within the whole species (Fig. 2b). To test the effect of the *PDR1*^*YJM326*^ allele in different backgrounds, we crossed the resistant isolate YJM326 with 20 diverse sensitive isolates. Counterintuitively, the resulting hybrids displayed continuous variation of the resistance in the presence of cycloheximide (Fig. 3a). To test whether the resistance variation in the hybrids were due to allelic interactions at the *PDR1* locus in different backgrounds, we introduced a plasmid carrying the *PDR1*^*YJM326*^ allele (pPDR1^YJM326^) into the same set of isolates, and quantified their fitness in the presence of cycloheximide (Fig. 3b). Across all isolates tested, about half (11/20) expressed the resistant phenotype to various degrees (Fig. 3b, Supplementary Fig. 6). However, fitness between haploid isolates carrying pPDR1^YJM326^ and the corresponding hybrids were only weakly correlated (Pearson’s correlation ρ = 0.434), indicating that allelic interactions at the *PDR1* locus only partly accounted for the observed variation (Supplementary Fig. 6).

**Fig 3.**
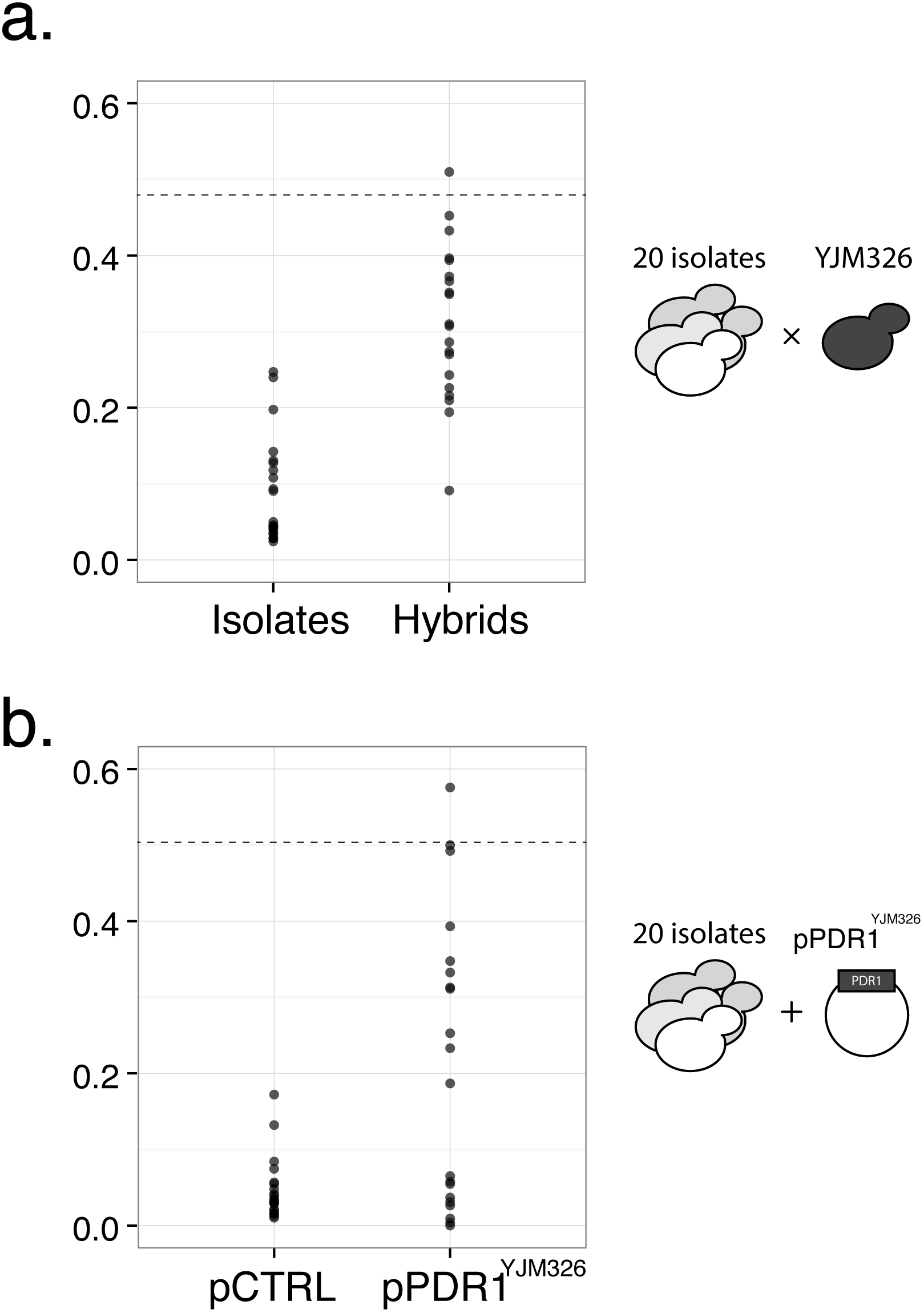
Effects of the *PDR1*^*YJM326*^ allele in different genetic backgrounds. (a) Fitness variation of 20 isolates (left panel) in comparison with the same set of strains hybridized with YJM326 in the presence of drug. Fitness values (y-axis) correspond to the ratio between the growth in the presence of cycloheximide (YPD CHX 1μg/ml) and control media YPD. Dashed line indicates the fitness of the resistant strain YJM326. (b) Fitness variation of 20 isolates carrying empty control plasmid (pCTRL, left panel) or plasmid containing the *PDR1*^*YJM326*^ allele under its native promoter (pPDR1^YJM326^, right panel). Fitness values were measured in the presence of cycloheximide (YPD CHX 1μg/ml) with hygromycin to maintain plasmid stability. Dashed line indicates the fitness value of YJM326 carrying the plasmid pPDR1^YJM326^. See Supplementary Figure 6 for detailed comparison for effect of hybrid and plasmid in individual genetic backgrounds.

The lack of correlation between hybrids and isolates carrying the plasmid with the *PDR1*^*YJM326*^ allele led us hypothesize the presence of potential modifiers in various hybrid backgrounds. To test this hypothesis, we evaluated the fitness distributions of the drug resistance in the offspring across the 20 hybrids generated previously. For each hybrid, 20 complete tetrads were tested in the presence of cycloheximide and the fitness distributions as well as the segregation patterns were assessed in the offspring (Supplementary Fig. 7). In the absence of modifiers, haploid segregants are expected to have complete phenotypic penetrance, as the effects of intralocus interaction were eliminated. In this scenario, all crosses between any sensitive parental isolate and YJM326 should display a bimodal distribution in the offspring, with a 2:2 segregation of the phenotype.

Interestingly, while most of the tested crosses (14/20) displayed Mendelian segregation as was observed in the cross between YJM326 and Σ1278b, several crosses showed clear deviation of the expected phenotypic distribution (Fig. 4, Supplementary Fig. 7). In addition to Mendelian cases (Fig. 4a), 3 other types of distribution were observed (Fig. 4b-d). In total, such cases represent ∼30% of all crosses tested (Fig. 4e). Of these crosses, 15% (3/20, between YJM320, Y3, Y9 and YJM326) showed incomplete penetrance, indicating possible suppressors of the *PDR1*^*YJM326*^ allele (Fig. 4b). We observed a 1:4:1 ratio between tetrads containing 2, 1 and 0 resistant segregants, possibly indicating that two independent loci, including *PDR1*, were involved (Fig. 4b, Supplementary Fig. 7). 10% of the crosses (2/20, between S288c, YJM440 and YJM326) showed enriched high fitness offspring, with an intermediate peak between the sensitive and resistant clusters. This observation suggests the presence of epistatic interactions from these specific genetic backgrounds, resulting as a transitional resistant phenotype cluster with higher genetic complexity (Fig. 4c). The levels of genetic complexity in these cases are suspected to be low, but the precise number of the genes involved remained unclear.

**Fig 4.**
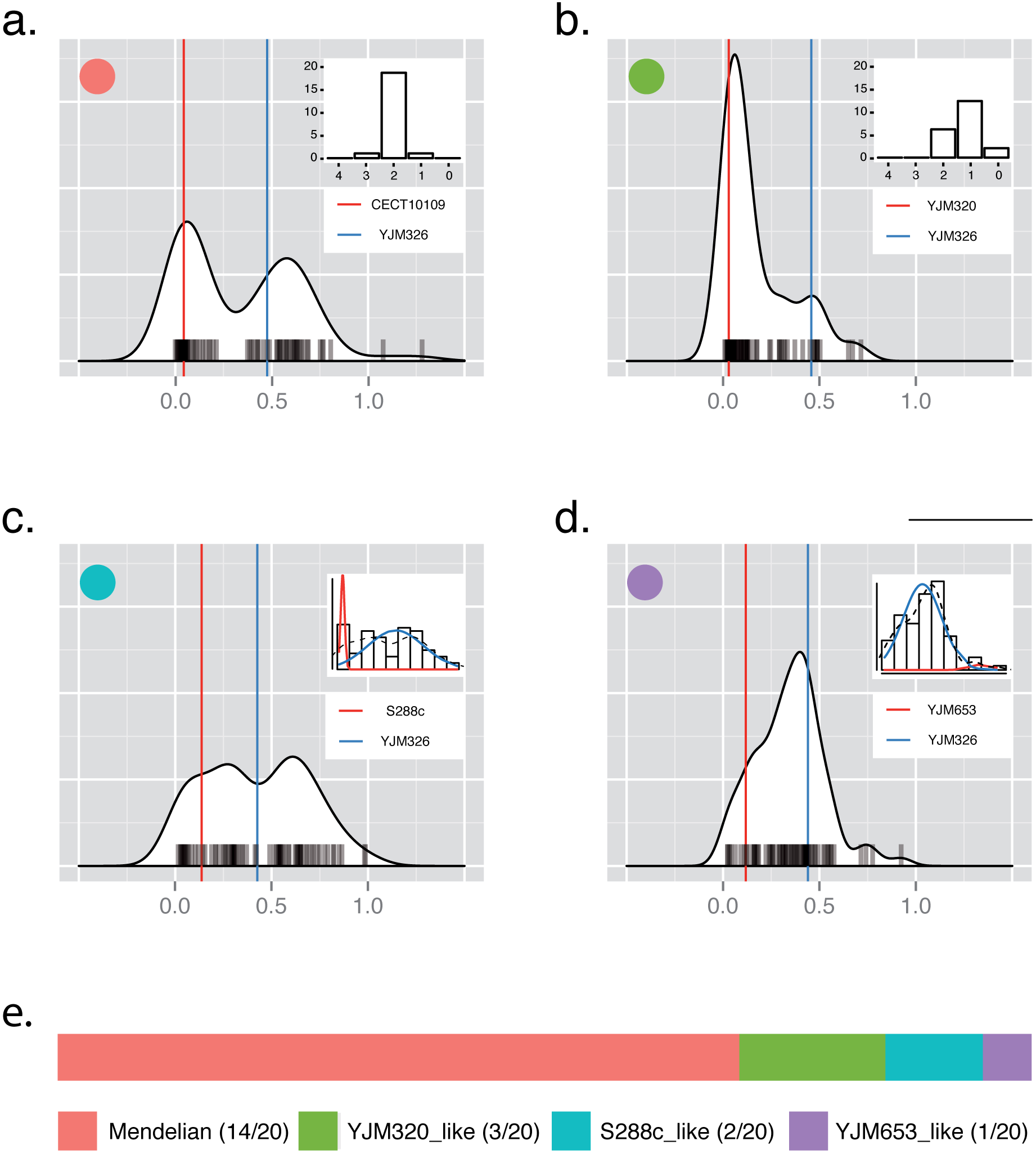
Post-Mendelian inheritance patterns of drug resistance in different hybrid contexts. (a-d) Offspring fitness distribution patterns observed in hybrids originated from 20 sensitive isolates and YJM326 in the presence of cycloheximide (YPD CHX 1μg/ml). 80 offspring were tested for each case, and examples of Mendelian (a) and non-Mendedian (c-d) inheritance patterns are shown. Phenotypic segregation is indicated at the upper right side. For non-bimodal cases the model fitting results were shown instead. Parental origins for each cross are shown, and the fitness values of the sensitive (red) or resistant (blue) parental strains are presented as vertical bars. (e) Distribution of different types of inheritance patterns observed. See Supplementary Figure 7 for offspring fitness distributions for each of the 20 crosses.

In addition to cases with low level of deviations from Mendelian expectations, we also found one cross (between YJM653 and YJM326) with a clear normal fitness distribution in the offspring. In this case, the resistant phenotype was no longer caused by a single Mendelian factor, and the underlying genetic determinants were undoubtedly complex (Fig. 4d). Contrasting to other identified Mendelian traits with a stable inheritance patterns across the population, the *PDR1* case represented a perfect example illustrating the hidden complexity of a simple Mendelian trait within natural population of the yeast *S. cerevisiae*.

## Discussion

By performing a species-wide survey of monogenic variants in *S. cerevisiae*, we obtained a first estimation of the proportion of Mendelian traits within a natural population. We showed that genes and alleles underlying the onset of Mendelian traits are variable in terms of their type, frequency and genomic distribution at the population level. Remarkably, by tracing the effect of one causal Mendelian variant *PDR1*^*YJM326*^ across the population, we demonstrated that the genetic complexity of traits could be dynamic, transitioning from clear Mendelian to diverse complex inheritance patterns depending on various genetic backgrounds.

Yeasts and more particularly *S. cerevisiae* have been extensively used as a model for dissecting many complex traits that were of medical, industrial and evolutionary interests^21-25^. A trend emerging from studying complex traits in this species was that causal variants do not distribute randomly across the genome, and several hotspots have been identified^26^. As a result, a low number of loci were found to be involved in high numbers of unrelated phenotypes, despite the fact that underlying causal genes could be different. Interestingly, causal variants in Mendelian traits seemed to follow the same trend as supported by our data. In fact, we observed phenotypic co-segregation of unrelated conditions such as resistance to acetate, 6-azauracil and osmotic stress, and showed that only a single region on chromosome IV was involved (Supplementary Fig. 4). In addition, the observed co-segregations showed relatively high population frequencies, with more than 15% of the crosses co-segregating on at least two different conditions (Fig. 1b). This effect of linkage could possibly lead to biased phenotype assortments across the population, although the underlying evolutionary origin is unknown.

In general, Mendelian traits were considered as rare especially in human disorders, however, no directly estimation of the proportion of Mendelian relative to complex traits was available at the population level, and what type of genes were more susceptible to cause Mendelian inheritance were unknown. Our data showed that across a yeast natural population, causal alleles involved in direct response to stress, such as transporters (*ENA*) or metal-binding genes (*CUP1*) were more likely to follow Mendelian inheritance. In fact, a large number of Mendelian traits identified in our sample were related to these two loci, and the inheritance patterns were extremely stable, displaying 2:2 segregations with little influence of the genetic backgrounds. Similar pattern was found in a Mendelian trait related ammonium resistance in natural isolates of *S. cerevisiae*, where a transporter gene *TRK1* was involved^27^. The stable inheritance patterns of traits caused by alleles with direct phenotypic effect could potentially due to the lack of regulatory complexity. As was supported by laboratory evolution experiments, amplifications of this type of genes were frequent, conferring to rapid acquisition of resistances in stress conditions such as salt^28^, copper^29,30^, sulfate^31^ and glucose limitations^32^.

By contrast, depending on the gene involved, a given Mendelian trait could lead to complex inheritance patterns across different genetic backgrounds, as evidence by the causal allele *PDR1* related to resistance to cycloheximide and anisomycin. By crossing the strain YJM326 carrying the resistant allele *PDR1*^*YJM326*^ with diverse natural isolates, we showed that although most crosses retained stable 2:2 segregations, the inheritance pattern of the resistance phenotype in some cases displayed various deviations from Mendelian expectation, including reduced penetrance (3/20), increased genetic complexity (2/20) and in one extreme case, transition from monogenic to complex trait. We propose that the observed post-Mendelian inheritance patterns are due to the functional nature of the *PDR1* gene. In fact, as *PDR1* encodes for a transcriptional factor with complex regulatory networks and impact multiple downstream effector genes^33^, the resulting phenotypic expression would possibly be influenced by variation of a large number of genes that are involved in the same network in different genetic backgrounds.

Overall, our data provided a first comprehensive view of natural genetic variants that lead to the onset of Mendelian traits in a yeast population. We showed that monogenic mutations could exhibit post-Mendelian modifications such as pleiotropy, incomplete dominance as well as variations in expressivity and penetrance due to differences in specific genetic backgrounds. Depending on the parental combination, the inheritance might display a Mendelian, intermediate or complex pattern, showing the continuum of the complexity spectrum related to a monogenic mutation, as illustrated by the example of the drug resistance involving *PDR1*^*YJM*^. However, while Mendelian traits could be related to common or rare variants, we found that the overall fitness distribution patterns of such traits at the population level, for some instances if not all, were not informative regarding their genetic complexity. Collectively, phenotypic prediction even for simple Mendelian variants may not be an easy task, in part due to the lack of prediction power using population data and the scarcity of large-scale family transmission information, such as the case for diseases in human. Future studies using pairwise crosses covering a larger panel of conditions in yeasts, or in other model organisms, may provide general trends and a more complete picture regarding the phenotypic predictability of monogenic traits.

## Materials and Methods

### Strains

Isolates from diverse ecological and geographical sources used in this study are detailed in Supplementary Table 1. All strains are stable haploids with deletion of the *HO* gene^34,35^. Laboratory strains FY4, FY5 (isogenic to S288c) and Σ1278b were used. Deletion mutants in the Σ1278b background were obtained from the gene deletion collection kindly provided by Charlie Boone^10^. YJM326 *Δpdr1* strain was generated by insertion of hygromycin resistance cassette *HygMX* using homologous recombination.

### Media and culture conditions

Detailed media compositions for phenotyping of the segregant panel are listed in Supplementary Table 2. Growth and maintenance of the strains are carried on standard rich media YPD (1% yeast extract, 2% peptone and 2% glucose). A final concentration of 200μg/ml hygromycin (Euromedex) was supplemented to maintain the plasmids carrying a resistance marker gene *HygMX*. Sporulation was induced on potassium acetate plates (1% potassium acetate, 2% agar). All procedures are performed at 30°C unless otherwise indicated.

### Crosses and generation of the offspring

For the construction of the segregant panel, 41 diverse isolates (*MAT*α) were crossed with the lab strain Σ1278b (*MAT*a) on YPD (Supplementary Table 1). Resulting diploids were sporulated for 2-3 days on sporulation medium (10 g/L potassium acetate, 20 g/L agar) at 30°C. Tetrad dissections were performed using the MSM 400 dissection microscope (Singer instrument) on YPD agar after digestion of the tetrad asci with zymolyase (MP Biomedicals MT ImmunO 20T). A total of 10 tetrads containing four viable spores were retained per cross. Same protocol was used for the validation of Mendelian cases with 2:2 segregation as well as the crosses between 20 isolates with the drug resistant strain YJM326 (Supplementary Table 1), in these cases 80 and 20 full tetrads were tested for each cross, respectively.

### High-throughput phenotyping and growth quantification

Quantitative phenotyping was performed using end point colony growth on solid media. Strains were pregrown in liquid YPD medium and pinned onto a solid YPD matrix plate to a 384 density format using a replicating robot RoTor (Singer instruments). At least two replicates of each parental strain were present on the corresponding matrix. The matrix plates were incubated overnight to allow sufficient growth, which were then replicated on 31 media conditions including YPD as a pinning control (see detailed compositions of the media in Supplementary Table 2). The plates were incubated for 48 hours at 30°C and were scanned at the 24, 40, 48 hour time points with a resolution of 600 dpi at 16-bit grayscale. Quantification of the colony size was performed using the Colony Area plugin in ImageJ, and the fitness of each strain on the corresponding condition was measured by calculating the normalized growth ratio between stress media and YPD using the software package ScreenMill^36^.

### Model fitting procedure and detection of traits with Mendelian inheritance

For each cross/trait combination, a bimodal distribution was fitted using the R package “mixtools” (https://cran.r-project.org/web/packages/mixtools/index.html) with k = 2 and maxit = 500. Mean (μ), standard deviation (σ), ratio between each cluster (λ) and posterior probability of each cluster for each individual were extracted from the output file. To determine cutoff values of posterior probability for cluster assignment, a simulated dataset was generated, by simulating two normal distributions with n*λ and n*(1-λ) individuals for each cluster, respectively, with mean and standard deviation randomly sampled from observations in real data. For each simulated set, the two normal distributions generated were combined, and the procedure was repeated for 1000 times to generate a training set with 1000 distributions (Supplementary Figure 1). The training set was then subjected to model fitting with the same parameters (Supplementary Figure 1). The mean (μ), standard deviation (σ), ratio between each cluster (λ) and posterior probability of each cluster for each individual were extracted again, and the training dataset was evaluated against the real data (Supplementary Figure 2). In this case, as the prior probability of cluster assignment was known for each simulated individual, it is possible to test for the detection sensitivity and specificity (Receiver Operating Characteristic or ROC) using varied cutoff parameters. A sequence starting from 0.5 to 0.95 (increment 0.05) for posterior probability and a sequence from 0 to 0.9 (increment 0.1) for percentage of non-overlapping individuals were tested. The ROC curves and area under the curve (AUC) were calculated for each combination of cutoff parameters using R package “ROCR”^37^. Cutoffs of 0.8 for posterior probability and 0.9 for percentage of non-overlapping were retained to ensure confident detections (Supplementary Figure 2). The defined parameters were applied on real data and cases passed the filter were preceded to cluster assignment (Supplementary Figure 1). For bimodal cases with parental pairs that belong to either phenotype cluster, the segregation patterns were determined. A trait is considered as Mendelian when more than 70% of the tetrads display a 2:2 segregation of the lethal phenotype. All analyses were performed in R.

### Bulk segregant analysis

In total, 6 crosses representative of the identified Mendelian traits were subjected to bulk segregant analyses in order to identify the genomic regions involved. Crosses involved were between Σ1278b and EM93, I14, YJM269 and CLIB272 in presence of salt and a second stressor, and between Σ1278b and YJM326 for antifungal drugs and copper sulfate. For each case, 50 independent viable spores tetrads exhibiting 2:2 segregation on the corresponding conditions were separately grown overnight at 30°C in liquid YPD and were pooled by equal optical density readings at 600nm. Pooled segregants were subjected to whole genome sequencing and the genomic regions involved in each trait were determined by looking at allele frequency variation.

### Genotyping strategy and data treatment

Genomic DNA from the pool was extracted using Genomic-tips 100/G columns and Genomic DNA buffers (QIAGEN) as described previously^38^. Sequencing of the samples was performed using Illumina Hiseq 2000 except for the cycloheximide pool, for which we used MiSeq technology. Reads were mapped to the Σ1278b genome with the Burrows-Wheeler Aligner (BWA, version 0.7.4) allowing 5 mismatches and 1 gap^39^. The ‘-I’ flag has been added for the MiSeq Pool because reads were encoded in Illumina 1.9 format. Single nucleotide polymorphism (SNP) calling has been performed using GATK v3.3-0^40^, with default parameters. The allele frequency of Σ1278b was calculated for each polymorphic position by adding the allele balance ratio, with the “VariantAnnotator” command of GATK.

### Reciprocal hemizygosity test

To perform reciprocal hemizygosity test on drug resistance, the wild type strains Σ1278b and YJM326 were crossed with each other and with deletion mutant strains of *PDR1* in both genetic backgrounds. Individual zygotes were isolated using the MSM 400 dissection microscope (Singer instrument) on YPD plate and the ploidy of the hybrids was checked on sporulation media after 2-3 days at 30°C. Phenotypic effects of the sensitive and resistance alleles were evaluated using drop test on selective media YPD CHX 1μg/ml and YPD as a growth control. Plates were scanned after 48 hours of incubation at 30°C.

### Plasmids construction and phenotyping

Centromeric plasmid was constructed to test the allelic effect of the drug resistance allele *PDR1*^*YJM326*^ in different genetic backgrounds using Gateway cloning technology (Invitrogen). Fragment containing *PDR1* and its native promoter and terminator regions flanked by attB1/attB2 recombination sites was amplified from the genomic DNA of YJM326 and Σ1278b, and was cloned into an empty centromeric plasmid with HphMX resistance marker pCTRL^22^, according to instructions. The resulting plasmids, pPDR1^YJM326^ and pPDR1^Σ1278b^ were verified using restriction enzymes and PCR amplification with internal primers of *PDR1* gene. 20 diverse natural isolates were transformed with pPDR1^YJM326^ as well as the empty control plasmid pCTRL using EZ transformation kit (MP biomedicals). Transformants were selected on YPD media containing 200 μg/ml hygromycin. Growth quantification of isolates carrying the pPDR1^YJM326^ or pCTRL plasmids in the presence of drug was performed as described previously, and 200 μg/ml hygromycin were supplemented to all media to maintain the selection pressure during the phenotyping procedure.

## Acknowledgments

The authors thank the National Institutes of Health (NIH Grant R01 GM101091-01) and the Agence Nationale de la Recherche (ANR Grant 2011-JSV6-004-01) for financial support. J.H. is supported in part by a grant from the Ministère de l’Enseignement Supérieur et de la Recherche and in part by a fellowship from the medical association La Ligue contre le Cancer.

## Accession numbers

All Illumina sequencing reads generated in this study have been deposited in the European Nucleotide Archive under the study accession number PRJEB11500.

